# TRPM7 activity drives human CD4 T-cell activation and differentiation in a magnesium dependent manner

**DOI:** 10.1101/2024.12.04.626765

**Authors:** Kilian Hoelting, Anna Madlmayr, Birgit Hoeger, Dorothea Lewitz, Marius Weng, Tanja Haider, Michelle Duggan, Rylee Ross, F. David Horgen, Markus Sperandio, Alexander Dietrich, Thomas Gudermann, Susanna Zierler

**Affiliations:** Walther Straub Institute of Pharmacology and Toxicology, Ludwig-Maximilians-Universität München, Goethestr.33, 80336 Munich, Germany; Institute of Pharmacology, Johannes Kepler University Linz, Altenbergerstr. 69, 4040 Linz, Austria; Laboratory of Marine Biological Chemistry, Hawai’i Pacific University, 1042 Fort Street Mall, Honolulu, HI 96813, USA; Institute of Cardiovascular Physiology and Pathophysiology, Biomedical Center, Ludwig-Maximilians-Universität München, Großharderner Str. 9, 82152 Planegg-Martinsried, Germany

## Abstract

T lymphocyte activation is a crucial process in the regulation of innate and adaptive immune responses. The ion channel-kinase TRPM7 has previously been implicated in cellular Mg^2+^ homeostasis, proliferation, and immune cell modulation. Here, we show that pharmacological and genetic silencing of TRPM7 leads to diminished human CD4 T-cell activation and proliferation following TCR mediated stimulation. In both primary human CD4 T cells and CRISPR/Cas-9 engineered Jurkat T cells, loss of TRPM7 led to altered Mg^2+^ homeostasis, Ca^2+^ signaling, reduced NFAT translocation, decreased IL-2 secretion and ultimately diminished proliferation and differentiation. While the activation of primary human CD4 T cells was dependent on TRPM7, polarization of naïve CD4 T cells into regulatory T cells (T_reg_) was not. Taken together, these results highlight TRPM7 as a key protein of cellular Mg^2+^ homeostasis and CD4 T-cell activation. Its role in lymphocyte activation suggests therapeutic potential for TRPM7 in numerous T-cell mediated diseases.

**Summary:** TRPM7 is crucial to maintaining cellular Mg^2+^ homeostasis and regulates human CD4 T-cell activation by modulating early Ca^2+^ signaling events in response to TCR-mediated stimulation subsequently, influencing T-cell differentiation in a Mg^2+^ dependent manner.

## Introduction

Immune cell function is essential for health and disease. Both innate and adaptive immune responses involve various cell types and are precisely regulated (Parenti et al., 2016; Walker, 2022). CD4 T lymphocytes are critically involved in both innate and adaptive immune responses (Parenti et al., 2016; Dong, 2021). Through different cellular subsets, CD4 T cells initiate adaptive immune responses against various kinds of pathogens. They have a crucial function in anti-cancer immunity, but also play a key role in the development of autoimmune diseases (Yatim & Lakkis, 2015; Bonilla & Oettgen, 2010; ABBAS, 2019; Walker, 2022). Robust receptor-mediated cell activation, including various costimulatory signals, is crucial for lymphocyte function and ultimately leads to cell proliferation and differentiation into specific effector cell types (Bonilla & Oettgen, 2010; Heinzel et al., 2018; Martínez-Méndez et al., 2021). Accordingly, T-cell activation is the target of several established and emergent pharmacological strategies for immune modulation. Thus, gaining further insights into T-cell activation and the involvement of interaction partners is necessary to gain a better understanding of potential therapeutic targets.

Melastatin-like Transient Receptor Potential, member 7 (TRPM7), is a protein ubiquitously expressed in mammals, showing high expression in lymphocytes (Beesetty et al., 2018; Krishnamoorthy et al., 2018). Embryonic development, thymopoiesis and cellular proliferation critically rely on TRPM7 activity (Beesetty et al., 2018; Nadler et al., 2001; Nadolni et al., 2020;). Expressing an ion channel in the plasma membrane, TRPM7 conducts divalent cations, such as Mg^2+^, Ca^2+^ and Zn^2+^ (Schmitz et al., 2003; Nadler et al., 2001; Liang et al., 2022). Mutations in the *TRPM7* gene are associated with several clinical phenotypes in humans and mice. Most of the symptoms induced by TRPM7-mediated pathologies including macrothrombocytopenia, reduced Mg^2+^ serum levels and signs of systemic inflammation, and can be by Mg^2+^ supplementation (Krishnamoorthy et al., 2018; Chubanov et al., 2024; Stritt et al., 2016; Sahni & Scharenberg, 2008). Different studies have characterized TRPM7 as a key player of cellular Mg^2+^ uptake (Cherepanova et al., 2016; Hoeger et al., 2023; Stritt et al., 2016), while other proteins proposed for this role, such as MagT1 transporter, have lost scientific support (Cherepanova et al., 2016; Li et al., 2011; Ravell et al., 2020). Moreover, the TRPM7 ion channel domain is covalently linked to a cytosolic serine/threonine kinase domain (Schmitz et al., 2003; Nadler et al., 2001; Liang et al., 2022). Different *in vitro* and native TRPM7 kinase substrates have been found, including myosin II, Annexin A1, phospholipase C gamma 2, SMAD2 and AKT (Clark et al., 2008; Dorovkov & Ryazanov, 2004; Romagnani et al., 2017; Hoeger et al., 2023). In recent years important insights have been gained regarding the role of TRPM7 in mammalian immune cells. Absence of TRPM7 channel function has been linked to reduced store-operated Ca^2+^ entry and proliferation arrest in DT40 chicken B cells and a kinase-deficient mouse model (Faouzi et al., 2017; Sahni & Scharenberg, 2008; Krishnamoorthy et al., 2018; Beesetty et al., 2018). Here, we shed light on the role of TRPM7 in human T lymphocyte homeostasis and activation. We demonstrated TRPM7 to be crucial for maintenance of cellular Mg^2+^ homeostasis, activation and proliferation of Jurkat T cells and primary CD4 T cells, as well as subsequent effector functions including cytokine release and polarization.

## Results

### TRPM7-mediated Mg^2+^ homeostasis is essential for Jurkat T-cell proliferation

Jurkat T cells are a well characterized and a commonly used cell line to study T lymphocyte function and signaling. We utilized this model to gain insights into the role of TRPM7 in functions of human T cells including T-cell activation. Applying CRISPR-Cas9 genome editing, we generated two clones of a novel TRPM7 KO Jurkat cell line harboring a genomic base pair insertion, which results in a frameshift in exon 4. The successful base pair insertion was confirmed through sequencing of the *TRPM7* gene (ThermoFisher). We were able to confirm the expected abolition of TRPM7 currents in these cells via whole cell patch-clamp experiments, thereby functionally verifying the knock-out (Fig. 1A, B and Suppl.Fig. 1A, B). While being morphologically indifferentiable to WT cells (data not shown), the cells of our TRPM7 KO clones showed a clear reduction of proliferation rates in standard Jurkat T cell media and died within five days. However, culturing these TRPM7 KO cells in media supplemented with 6 mM MgCl_2_ restored normal proliferation and prevented cell death (Fig. 1C, D and Suppl. Fig. 1C, D). To further examine the nature of the TRPM7 KO T cells’ need for MgCl_2_ supplementation, we performed inductively coupled plasma mass spectrometry (ICP-MS), which revealed a reduction of cellular magnesium content in TRPM7 KO cells (Fig. 1E and Suppl.Fig. 1E), while culturing them in medium supplemented with 6 mM MgCl_2_ restored intracellular Mg^2+^ levels (Fig. 1E and Suppl.Fig. 1E). In parallel, we employed the known pharmacological inhibitor of the TRPM7 channel, NS8593 (Chubanov et al., 2012), which similarly abolished TRPM7 currents in WT Jurkat T cells (Fig. 1F, G). Culturing WT Jurkat T cells in the presence of NS8593 produced a similar effect as the TRPM7 KO. Treatment markedly reduced cell proliferation and viability within five days, with survival and proliferation being partially restored by supplementing extracellular MgCl_2_ (Fig. 1H, I). Since NS8593 has been known to also inhibit SK2-channels in other cell types, we controlled for a potential SK2-dependent effect by employing the SK2-inhibitor apamin, which did not influence TRPM7 currents in respective patch-clamp experiments (Suppl. Fig. 2 A, B). Apamin likewise did not affect lymphocyte growth and viability (Suppl. Fig. 2 C, D). Similar to Jurkat TRPM7 KO clones, treatment with NS8593 also resulted in reduced cellular Mg^2+^ levels, as analyzed by ICP-MS (Fig. 1J). Likewise, Mg^2+^ supplementation of the medium restored intracellular Mg^2+^ levels (Fig. 1J). In line with previous studies on TRPM7 (Zierler et al., 2011), these findings emphasize the importance of the channel for cell proliferation and Mg^2+^ homeostasis in Jurkat T cells.

**Figure 1:**
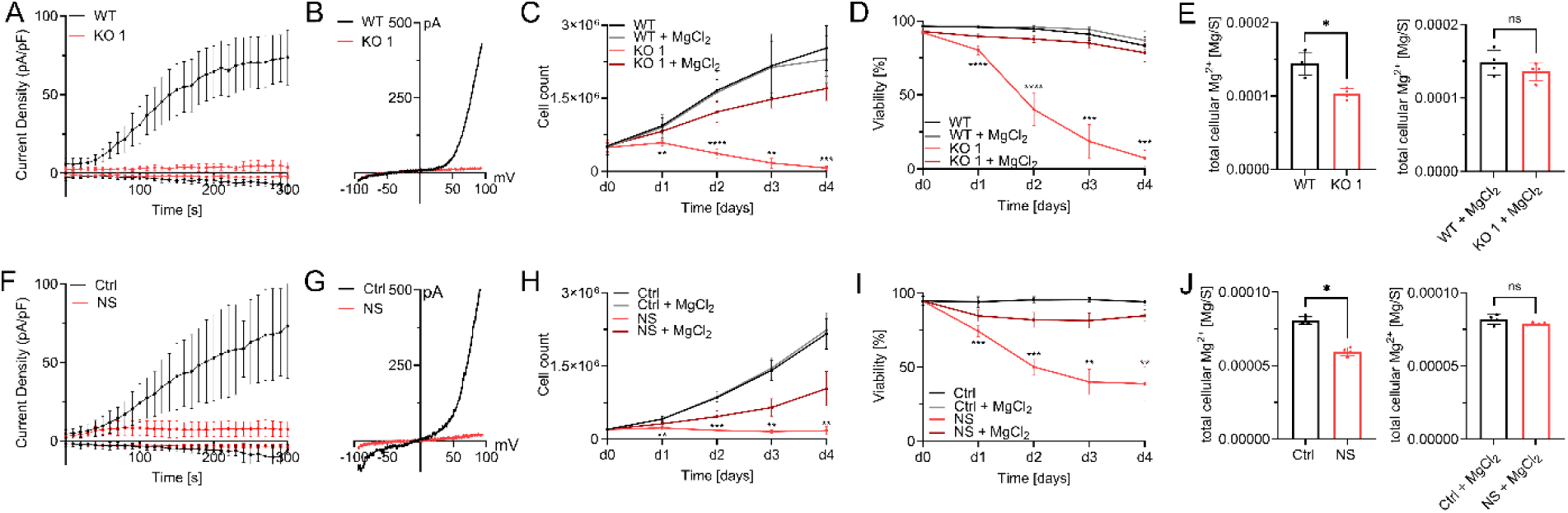
TRPM7-mediated Mg^2+^ homeostasis is essential for Jurkat T-cell proliferation. A) TRPM7 current densities and B) TRPM7 I/V relationship of Jurkat cells during whole-cell patch clamp experiment with Mg^2+^-free intracellular solution. WT (WT, black) and TRPM7 KO Jurkat clones (KO, red), n(WT)=9; n(KO)=10. C) Cell counts and D) viability of natively proliferating TRPM7 WT and KO Jurkat clones in RPMI medium with 10% FBS, with and without supplementation with 6 mM MgCl_2_, n=3, measured in duplicates. E) Cellular Mg^2+^ content quantified by ICP-MS. WT and TRPM7 KO Jurkat clones, cultured in regular (WT-)media for 18 h ahead of sampling, n=4. And WT and TRPM7 KO Jurkat clones, cultured in regular (WT-)media supplemented with 6 mM MgCl_2_ for 18 h ahead of sampling, n=4. F) TRPM7 current densities and G) TRPM7 I/V relationship of Jurkat cells during whole-cell patch clamp with Mg^2+^-free intracellular solution. WT Jurkat cells, treated with DMSO as solvent control (Ctrl, black) or treated with 30 µM NS8593 (NS, red), n(Ctrl)=6; n(NS)=10. H) Cell counts and I) viability of natively proliferating Jurkat cells in RPMI medium with 10% FBS, with and without supplementation with 6 mM MgCl_2_, and treated with DMSO as solvent control (Ctrl, black) or treated with 30 µM NS8593 (NS, red), n=4. J) Cellular Mg^2+^ content as measured with ICP-MS. Jurkat WT cells, treated with DMSO as solvent control (Ctrl, black) or treated with 30 µM NS8593 in DMSO (NS, red), cultured in regular (WT-) media without and with supplementation with 6 mM MgCl_2_ for 18 h ahead of sampling, n=4. Statistics: Two-way ANOVA (C, D, H, I) or one-way ANOVA (E, J). * P<0.05; ** P<0.005; *** P<0.0005 and **** P<0.0001. Data are mean ± SD.

### TRPM7 channel activity is essential for Jurkat T-cell activation

Having tested the general functionality of our genetic and pharmacological models in Jurkat T cells, we proceeded with studies to decipher the role of TRPM7 in the activation process of human lymphocytes. Previously, TRPM7 was linked to altered store-operated Ca^2+^ entry (SOCE) in DT40 chicken B lymphocytes (Faouzi et al., 2017). As an important early step in lymphocyte activation, we designed our experiments to first characterize the effects of TRPM7 in Ca^2+^ signaling. Using Fura-2 as a ratiometric Ca^2+^ indicator, we performed Ca^2+^ imaging experiments comparing Jurkat TRPM7 WT and KO cells. Following depletion of the intracellular Ca^2+^ stores using thapsigargin, TRPM7 KO cells exhibited a strongly reduced rise in cytosolic Ca^2+^ concentration ([Ca^2+^]_i_) (Fig. 2A and Suppl. Fig. 1F), suggesting SOCE to be defective in Jurkat T cells lacking TRPM7. We performed the experiment with Jurkat T cells in the absence and presence of the specific TRPM7 channel inhibitor NS8593. Similar to the effect seen in the KO model, cells treated with the blocker exhibited a strong reduction of the [Ca^2+^]_i_ elevation (Fig. 2G). To quantify the amount of Ca^2+^ present in the cytosol during the measurement, we calculated the area under the curve of the Ca^2+^ traces (Fig.2B and 2H respectively and Suppl. Fig. 1G). They, too, show a marked reduction of [Ca^2+^]_i_ in both the KO T cells and the NS8593 treated Jurkat T cells, indicating an early activation defect. This Ca^2+^ signaling defect would likely affect subsequent transcription factor recruitment. Given that an increase in [Ca^2+^]_i_ is directly responsible for calcineurin-mediated dephosphorylation and subsequent nuclear translocation of NFAT molecules (Maguire et al., 2013; Park et al., 2020; Lin et al., 2019), we next tested Ca^2+^ induced NFATc1 translocation. Basal levels of nuclear NFATc1 were comparable in WT and KO cells. Again, using thapsigargin as stimulant, we were able to induce the translocation of NFATc1 to the nucleus in WT control cells. Thapsigargin-induced translocation was diminished in both in TRPM7 KO cells and in cells treated with NS8593 (Fig. 2C-D and I-J respectively and Suppl. Fig. 1H-I). Having observed altered transcription factor recruitment, we assessed mRNA expression levels of *IL-2,* a well-known NFAT target gene (Maguire et al., 2013; Sakellariou et al., 2024). Both, TRPM7 KO cells and cells after application of the TRPM7 inhibitor showed a remarkable reduction of *IL-2* mRNA (Fig. 2E, K respectively). One important feature of T-cell activation is the expression of activation markers on the cell surface, of which CD69 is robustly upregulated in stimulated Jurkat T cells. In line with data shown by Mendu et al., who found an upregulation of CD69 in TRPM7-deficient murine thymocytes (Mendu et al., 2020), representative FACS plots for gating strategy are shown in Suppl. Fig. 3A, depicted a similar picture for human Jurkat T cells. 24 h after activation, viable TRPM7 WT and KO cells upregulated CD69 to a similar extent (Fig. 2F). Interestingly, treatment with NS8593 lead to a significant reduction of CD69 upregulation in Jurkat T cells, (Fig. 2M), while apamin treatment did not affect CD69 upregulation (Suppl. Fig. 2E). Thus, treatment with a TRPM7 blocker affected T-cell activation whereas genetic TRPM7 ablation did, possibly because TRPM7 KO cells had developed compensatory mechanisms, in clear contrast to the acute blockade of TRPM7 activity by its specific inhibitor. Overall, these data show a role of TRPM7 in modulating Ca^2+^ signaling and downstream Ca^2+^ dependent translocation of transcription factors and gene expression.

**Figure 2:**
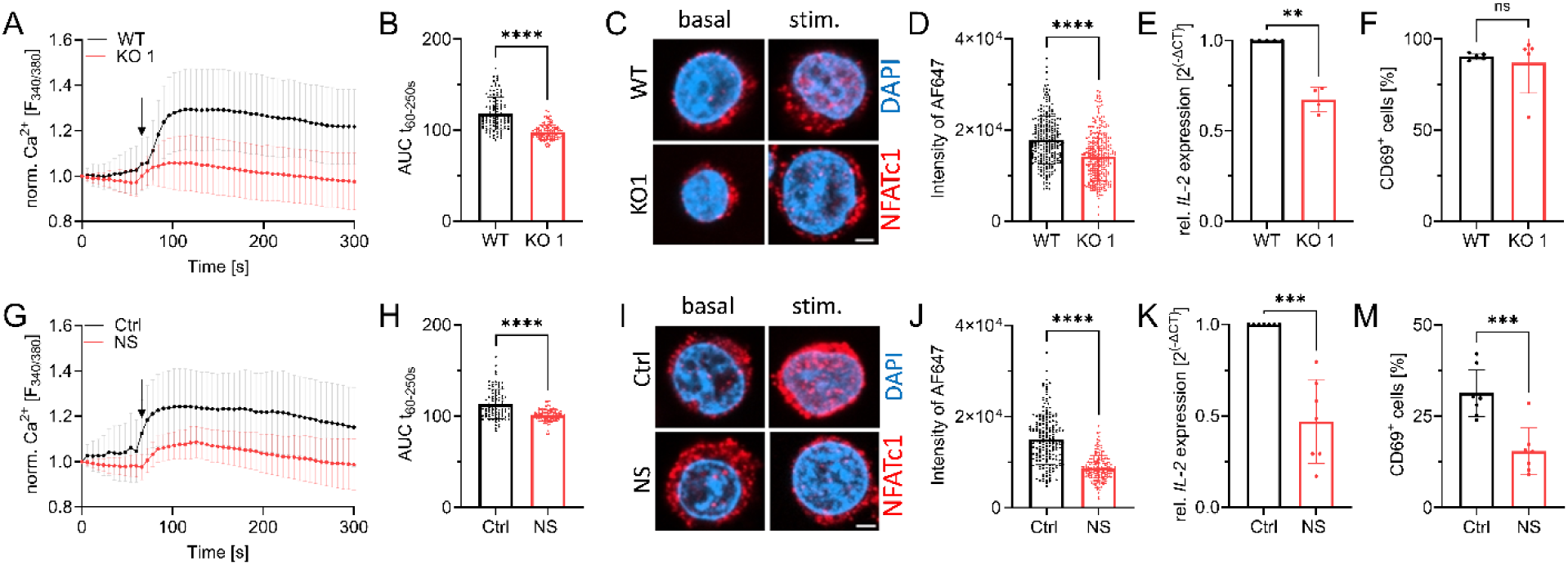
TRPM7 is essential for Jurkat T-cell activation. A) Fura-2 based imaging of cytosolic Ca^2+^ concentration of Jurkat cells. Stimulation with 5 µM thapsigargin at the indicated time point (arrow) of WT (WT, black) and TRPM7 KO (KO, red) Jurkat cells, n(WT)=111; n(KO)=113. B) Quantification of the area under the curve (AUC) of respective curves shown in A. C) Representative immune-fluorescent images of the NFATc1 localization in WT and KO cells before (basal) and after 30 min stimulation (stim.) with 5 µM thapsigargin, scale bar = 2 μm. NFATc1 in red, DAPI in blue. D) Quantification of nuclear NFATc1 levels (corresponding to AF647 signal intensity) upon stimulation of TRPM7 WT (WT, black) and KO (KO, red) cells, n(WT)= 261; n(KO)=279. E) Relative *IL-2* mRNA expression levels of Jurkat WT (WT, black) and KO (Ko, red) cells, n=4. F) CD69 expression of stimulated Jurkat cells, WT (WT, black) and KO (KO, red), n=5. G) Ca^2+^ imaging of WT Jurkat, treated with DMSO as solvent control cells (Ctrl, black) or cells treated with 30 µM NS8593 (NS, red). Stimulation with 5 µM thapsigargin at indicated time point (arrow), n(Ctrl)=95; n(NS)=94. H) Quantification of the area under the curve (AUC) of respective curves shown in G. I) Representative immune-fluorescent images of NFATc1 localization in DMSO treated cells as solvent control (Ctrl, black) or treated cells with 30 µM NS8593 (NS, red) before and after 30 min stimulation with 5 µM thapsigargin, scale bar = 2 μm. J) Quantification of nuclear NFATc1 levels upon stimulation of cells treated with DMSO as solvent control (Ctrl, black) or cells treated with 30 µM NS8593 (NS, red), n(Ctrl)=196; n(NS)=195. K) Relative *IL-2* mRNA expression levels of cells treated with DMSO as solvent control (Ctrl, black) or cells treated with 30 µM NS8593 (NS, red), n=7. M) CD69 expression of cells treated with DMSO as solvent control (Ctrl, black) or cells treated with 30 µM NS8593 (NS, red), after α-CD3 stimulated, n=6-7. Statistics: Student’s t test (B, D, F, H, I, M) and Mann-Whitney U test (E, K). ** P<0.005; *** P<0.0005; **** P<0.0001 and n.s.—not significant. Data are mean ± SD.

### TRPM7 inhibition alters Ca^2+^ signaling and NFAT translocation in primary human CD4 T lymphocytes

Having validated NS8593 as an applicable pharmacological tool able to mimic the absence of TRPM7 protein in lymphocytes, we broadened the scope of the study to primary human CD4 T cells. Studying primary human lymphocytes instead of cell lines strongly increases the transferability of *in vitro* findings to immunological processes in human health and disease. CD4 T lymphocytes, isolated from healthy human PBMCs, were used to shed light on both naïve as well as conventional (CD4^+^ CD25^-^ effector) CD4 T cells. Isolated populations were validated by Flow Cytometry (Suppl. Fig. 3B, C). Using whole-cell patch clamp, we were able to show functional channel expression of TRPM7 in naïve CD4 T cells and the conventional CD4 T cell population. In both cell populations TRPM7 currents were absent after treatment with NS8593 (Fig. 3A, G). Analogous to our Jurkat experiments, we characterized the Ca^2+^ dependent activation cascade of primary CD4 T cells. We used antibodies against CD3 and CD28 to elicit TCR-dependent Ca^2+^ signaling, which was analyzed by Fura-2 based Ca^2+^ imaging. After applying stimulating antibodies to isolated naïve primary human CD4 T cells, a robust increase in [Ca^2+^]_i_ followed by oscillations of Ca^2+^ concentration, in a large subset of T cells (Fig. 3B). Cells treated with the specific TRPM7 channel inhibitor NS8593 showed no reduction in basal Ca^2+^ influx as well as in changes in intracellular Ca^2+^ concentrations (Fig. 3C-E), but had altered kinetics of [Ca^2+^]_i_ increase. Importantly, cytosolic Ca^2+^ oscillations, which have been shown to be crucial for activation-induced gene expression, were absent upon TRPM7 inhibition (Fig. 3F). Studying the CD4^+^ CD25^-^ effector T cell population, also referred to as conventional CD4 T lymphocytes, displayed similar results. The average Ca^2+^ concentration increased similarly, but showed altered kinetics. NS8593, as a specific TRPM7 inhibitor, almost eliminated Ca^2+^ oscillations in treated cells (Fig. 3H-L). Application of the SK2 channel inhibitor apamin, however, did not reduced Ca^2+^ oscillations (Suppl. Fig. 2F). With both the amount of Ca^2+^ as well as the characteristic Ca^2+^ oscillations known to be crucial for NFAT translocation to the nucleus (Maguire et al., 2013; Park et al., 2020; Lin et al., 2019), we proceeded by studying this process. We quantified NFATc1 residing in the nucleus after TCR-mediated stimulation in naïve and conventional CD4 T cells, as well as in cells treated with NS8593. Here, we saw in both cell subsets that TRPM7 inhibition resulted in reduced activation-dependent NFAT-translocation (Fig. 3M-P). This NS8593-induced defect in NFATc1-translocation highlights the importance of the Ca^2+^-oscillations, which were also diminished in cells with TRPM7 blockade (Fig. 3M-P). These results suggest an important role of TRPM7 in the early activation process of primary naïve and conventional CD4 T cells with large implications on activation-dependent gene expression.

**Figure 3:**
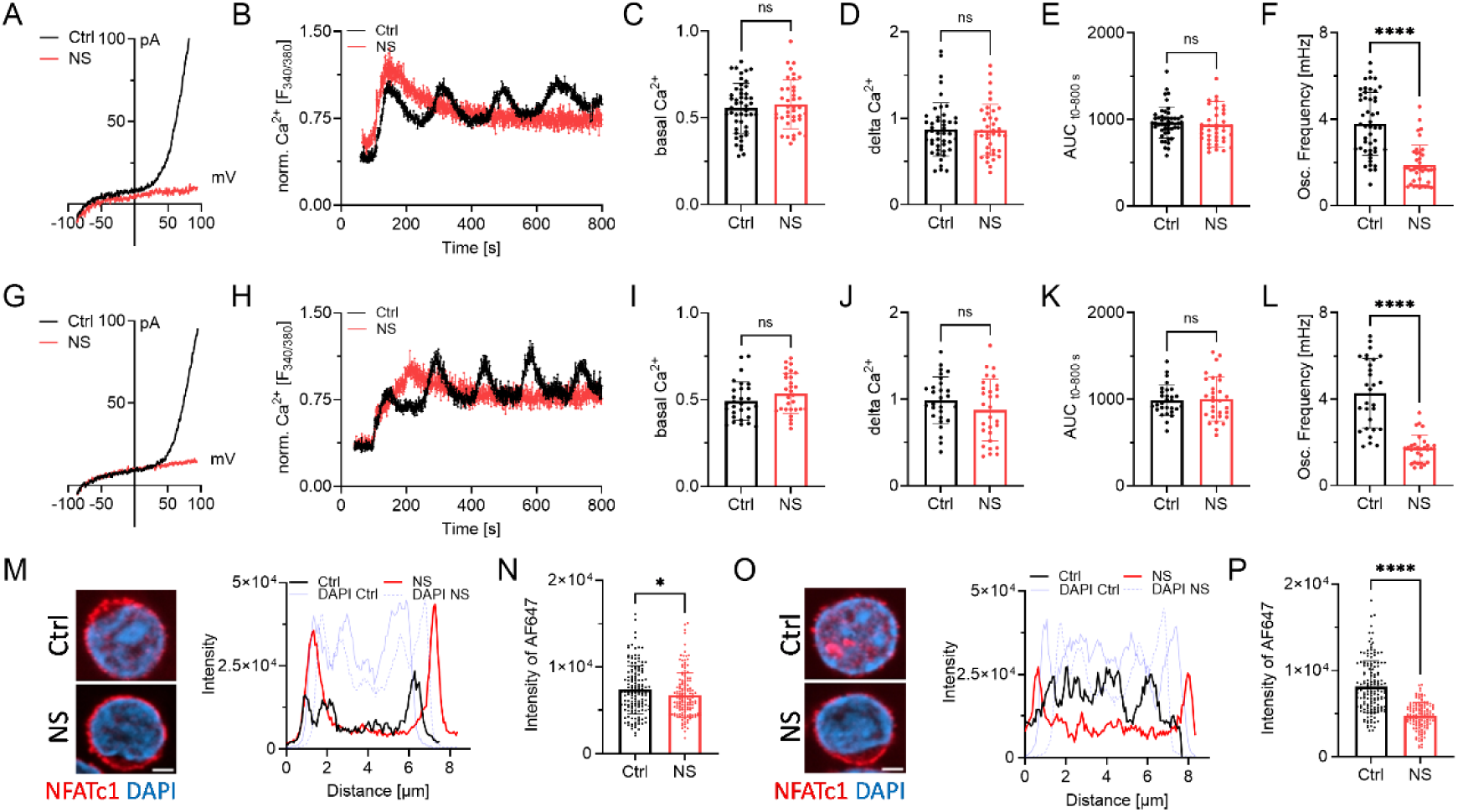
TRPM7 inhibition alters Ca^2+^ signaling and NFAT translocation in primary human CD4 T lymphocyte. A) TRPM7 I/V relationship of naïve CD4 T cells during whole-cell patch clamp with Mg^2+^-free intracellular solution. Cells treated with DMSO as solvent control (Ctrl, black) or cells treated with 30 µM NS8593 (NS, red). B) Representative trace of naïve CD4 T cells Fura-2 based imaging of cytosolic Ca^2+^ concentrations following anti-CD3/CD28 stimulation. Antibodies bound to microscopy chamber bottom with cells sinking down in saline containing 2 mM Ca^2+^ during running measurement, coming to rest in focus plane with contact to stimulation antibodies. Cells treated with DMSO as solvent control (Ctrl, black) or treated with 30 µM NS8593 (NS, red). Respective quantification of Ca^2+^ imaging experiments of naïve CD4 T cells for C) basal, D) delta Ca^2+^, E) AUC and F) oscillation frequency, n=29-30 cells. G) TRPM7 I/V relationship of conventional CD4 T cells during whole-cell patch clamp with Mg^2+^-free intracellular solution. Cells treated with DMSO as solvent control (Ctrl, black) or cells treated with 30 µM NS8593 (NS, red), n(Ctrl)=5, n(NS)=5. H) Representative trace of conventional CD4 T cells Fura-2 based imaging of cytosolic Ca^2+^ concentrations following anti-CD3/CD28 stimulation. Antibodies bound to microscopy chamber bottom with cells sinking down in saline containing 2 mM Ca^2+^ during running measurement, coming to rest in focus plane with contact to stimulation antibodies. Cells treated with DMSO as solvent control (Ctrl, black) or treated with 30 µM NS8593 (NS, red). Respective quantification of Ca^2+^ imaging experiments of conventional CD4 T cells for I) basal, J) delta Ca^2+^, K) AUC and L) oscillation frequency, n= 39-48 cells. M) Representative immune-fluorescent images of NFATc1 localization (NFATc1 in red, DAPI in blue) and intensity profile of subcellular NFATc1 distribution (Ctrl in black, NS in red, respective DAPI in blue) of naïve CD4 T cells treated with DMSO as solvent control and TRPM7 inhibited cells upon 30 min stimulation with anti-CD3/CD28, scale bar = 2 μm. N) Quantification of nuclear NFATc1 levels upon stimulation of cells treated with DMSO as solvent (Ctrl, black) or in presence of 30 µM NS8593 (NS, red) cells, n(Ctrl)=149; n(NS)=144. O) Representative immune-fluorescent images of NFATc1 localization (NFATc1 in red, DAPI in blue) and intensity profile of subcellular NFATc1 distribution (Ctrl in black, NS in red, respective DAPI in blue) of conventional CD4 T cells of Ctrl and TRPM7-inhibited cells upon 30 min stimulation with anti-CD3/CD28, scale bar = 2 μm. NFATc1 in red, DAPI in blue. P) Quantification of nuclear NFATc1 levels upon stimulation and treatment with DMSO as solvent control (Ctrl, black) or in presence of 30 µM NS8593 (NS, red) cells, n(Ctrl)=155; n(NS)=132. Statistics: Student’s t test (C-F, I-L, N, P). * P<0.05; **** P<0.0001 and n.s.—not significant. Data are mean ± SD.

### TRPM7 inhibition affects activation of primary human CD4 T cells

As transcription factor recruitment is crucial for *IL-2* expression (Maguire et al., 2013; Sakellariou et al., 2024), we next investigated the stimulation-dependent release of this autocrine and paracrine cytokine of CD4 T cells. After 48 h stimulation control cells had secreted significantly more *IL-2* into the supernatant than cells treated with NS8593. This effect could be partially rescued by MgCl_2_ supplementation (Fig. 4A, F). We next investigated activation-induced protein expression. Upregulation of CD69 and CD25 are important hallmarks of T-cell activation, both being physiologically significant and well-studied (Nisnboym et al., 2023; Peng et al., 2023; Poloni et al., 2023). In response to CD3/CD28-stimulation, both activation markers were upregulated in primary CD4 lymphocyte cells, shown by representative FACS plots and gating strategy in Suppl. Fig. 3A. Both in naïve CD4 T cells (Fig. 4B-E) and conventional CD4 T cells (Fig. 4G-J) treated with NS8593, upregulation of CD69 and CD25 was markedly reduced, an effect that could be reverted with MgCl_2_ supplementation. MgCl_2_ supplementation also increased the upregulation of activation marker in control cells, underlining the importance of Mg^2+^ in T-cell activation (Fig. 4B-E and G-J). While TCR-mediated CD69- and CD25-upregulation was, as expected, less pronounced in naïve T cells compared to the conventional CD4 T cells, inhibition of TRPM7 yielded similar effects in both cell populations (Fig. 4B-E and G-J). Titration of inhibitor NS8593 showed a dose-dependent reduction of CD69 and CD25 upregulation in CD4 T cells (Suppl. Fig. 4B, C). To improve methodic robustness, we repeated our experiments with another known specific TRPM7 channel inhibitor, waixenicin A (Zierler et al., 2011). By whole-cell patch clamp, we were able to confirm blockade of TRPM7 currents upon pharmacological treatment with waixenicin A (Fig. 4K). Both inhibitors yielded a very similar upregulation of CD69 and CD25 in these cells upon TCR-mediated stimulation (Fig. 4L-O), which strongly supports a TRPM7-dependent effect. In summary, TRPM7 to affects transcription marker recruitment, IL-2 secretion and the upregulation of activation-dependent surface markers in both, naïve and conventional CD4 T cells.

**Figure 4:**
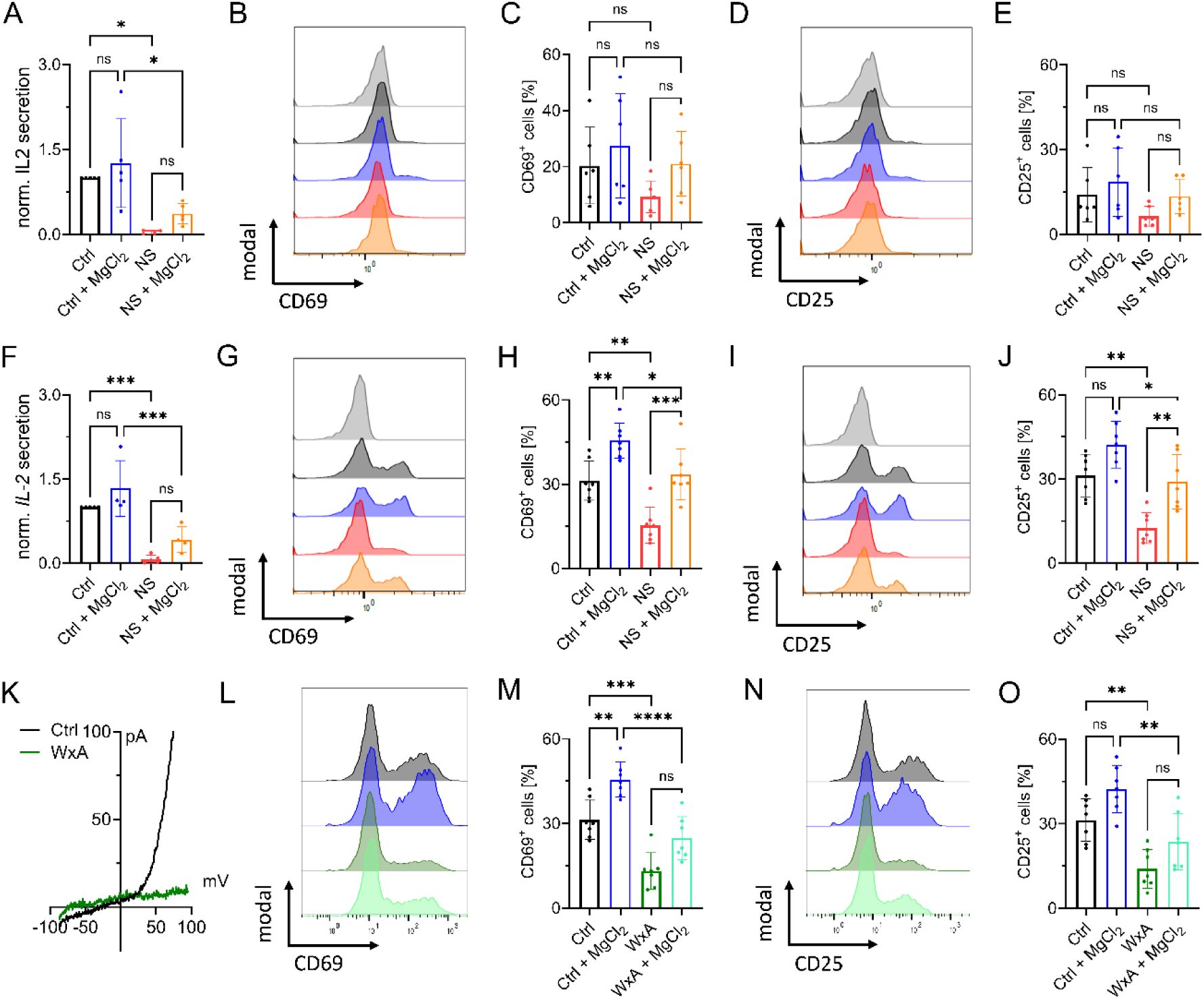
TRPM7 inhibition affects activation of primary human CD4 T cells. A) *IL-2* quantification of supernatant of naïve CD4 T cells 48 h after anti-CD3/CD28 stimulation, n=4-5. Histograms and quantification of upregulated activation markers CD69 (B-C) and CD25 (D-E) in naïve CD4 T lymphocytes 48 h after stimulation. Cells treated either with DMSO as solvent control (Ctrl, black) or with 30 μM NS8593 (NS, red), both with and without supplementation with 6 mM MgCl_2_. F) *IL-2* quantification of supernatant of conventional CD4 T cells 48 h after anti-CD3/CD28 stimulation or cells treated with DMSO as solvent control (Ctrl, black) or with 30 µM NS8593 (NS, red), both with and without supplementation with 6 mM MgCl_2_, n=4-5. Histograms and quantification of upregulated activation markers CD69 (G-H) and CD25 (I-J) in conventional CD4 T lymphocytes 48 h after stimulation. Cells treated either with DMSO as solvent control (Ctrl, black) or 30 μM NS8593 (NS, red), both with and without supplementation with 6 mM MgCl_2_. K) TRPM7 I/V relationship of conventional CD4 T cells during whole-cell patch clamp with Mg^2+^-free intracellular solution. Cells treated with EtOH as solvent control (Ctrl, black) or cells treated with 10 µM waixenicinA (WxA, green). Histograms and quantification of upregulated activation markers CD69 (L-M) and CD25 (N-O) in conventional CD4 T lymphocytes 48 h after stimulation. Cells treated either with EtOH as solvent control (Ctrl, black) or 10 μM waixenicinA (WxA, green), both with and without supplementation of 6 mM MgCl_2_, n=7. Statistics: One-way ANOVA (A, C, E, F, H, J, M, O). * P<0.05; ** P<0.005; *** P<0.0005; **** P<0.0001 and n.s.—not significant. Data are mean ± SD.

### TRPM7-induced Mg^2+^ deficiency promotes human naïve CD4 T cell to iT_reg_ differentiation

In proliferation experiments following anti-CD3/CD28 stimulation, we observed robust proliferation of the activated CD4 control cells within five days. Treatment with NS8593 strongly reduced cell proliferation (Fig. 5A, B). This effect was dose-dependent and could be partially reversed by supplementation with MgCl_2_ (Fig. 5A, B). An important hallmark of adaptive immunity and a consequence of successful T-cell activation is increased proliferation, clonal expansion and differentiation. Mendu et. al recently linked TRPM7 with thymic development of regulatory T cells (T_reg_) cells in a TRPM7 knockout mouse model (Mendu et al., 2020). Thus, we investigated the role of TRPM7 in the differentiation of naïve CD4 T cells to iT_regs_. Interestingly, in the presence of the TRPM7 inhibitor NS8593, we observed a reduction of CD25^+^ iT_regs_ (Fig 5C), correlating with our data on reduced CD4 T-cell activation upon TRPM7 inhibition. However, the successfully differentiated cells showed a higher FOXP3 expression upon NS8593 treatment compared to control (Fig. 5D-E). Repeating these experiments with the afore employed specific TRPM7 inhibitor waixenicin A, showed similar results. In addition, our experiments revealed a negative effect of Mg^2+^ on T_reg_ polarization, which could be rescued with TRPM7 inhibition (Fig. 5F, G). These findings point towards a modulatory role of TRPM7 in iT_reg_ differentiation, most likely by controlling Mg^2+^ homeostasis, as summarized in Fig. 5H.

**Figure 5:**
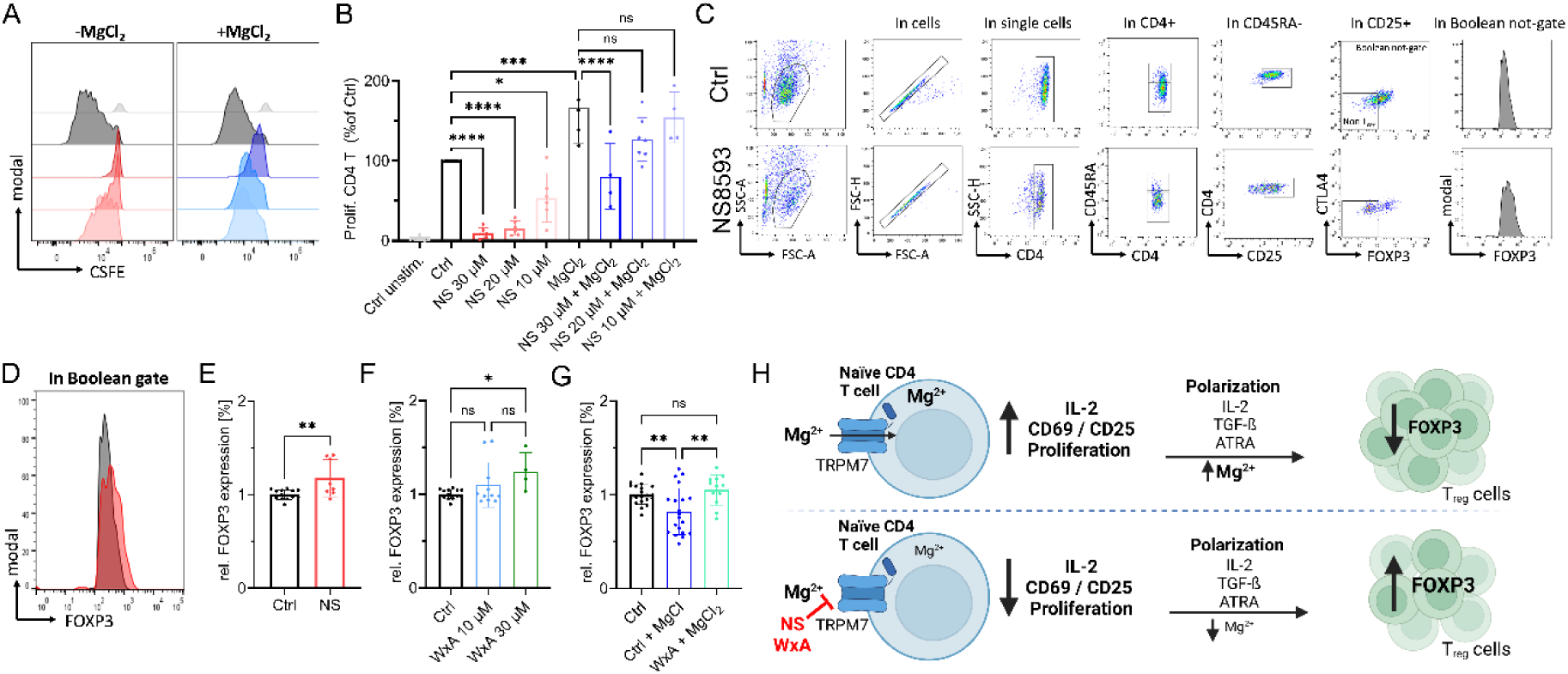
TRPM7-induced Mg^2+^ deficiency promotes human naïve CD4 T cell iT_reg_ differentiation. A) Representative histograms of dose-dependent proliferation (CSFE dye dilution) of conventional CD4 T cells in presence of various NS8593 concentrations, with (right) and without (left) supplementation with 6 mM MgCl_2_. Cells gated on T cell population, single cells and CD4 cells. Color code as in B. Cells gated on T cell population, single cells and CD4 T cells. B) Respective quantification of NS8593 dose dependent proliferation of conventional CD4 T cells, with and without supplementation with 6 mM MgCl_2_, corresponding to H, n=4-7. C) Representative FACS plots and gating path of iTreg cells after 6 days of differentiation of naïve CD4 T cells, cells treated with DMSO as solvent control (upper panel) or treated with 30 µM NS8593 (lower panel). D) Representative histogram overlay of FOXP3 signal in Boolean gate of DMSO controls (Ctrl, black) or in presence of 30 µM NS8593 (NS, red). E) Respective quantification of FOXP3 signal of cells treated with DMSO as solvent control (Ctrl, black) or 30 µM NS8593 (NS, red), n(Ctrl)=14; n(NS)=8. F) Respective quantification of FOXP3 signal of cells treated with EtOH as solvent control (Ctrl, black), 10 µM waixenicin A (WxA, blue) or 10 µM waixenicin A (WxA, green), n(Ctrl)=14; n(10 µM WxA)=11; n(30 µM WxA)=4. G) Respective quantification of FOXP3 signal of EtOH controls (Ctrl, black), DMSO Ctrl + 6 mM MgCl_2_ (Ctrl+MgCl_2_, blue), 30 µM waixenicin A + MgCl_2_ (WxA+MgCl_2_, turquoise), n(Ctrl)=20; n(Ctrl+MgCl_2_)=20; n(30 µM WxA + MgCl_2_)=12. H) Graphical summary of TRPM7-independent iT_reg_ differentiation. Pharmacological blockade of TRPM7 reduces intracellular Mg^2+^ levels and results in reduced IL-2 secretion, impaired upregulation of T-cell activation markers CD69 and CD25 and diminished proliferation in presence of TCR stimulus. TRPM7 inhibition followed by polarization of naïve CD4 T cells in presence of anti-CD3/CD28, IL-2, TGF-ß and ATRA, an iT_reg_ polarization cocktail, results in lower iT_reg_ numbers but enhanced FOXP3 expression. Figure created in https://BioRender.com. Statistics: One-way ANOVA (B, F, G) and Student’s t test (E). * P<0.05; ** P<0.005; *** P<0.0005; **** P<0.0001 and n.s.—not significant. Data are mean ± SD.

Altogether, our collective results depict TRPM7 as a primary player of T-cell activation and cellular Mg^2+^ homeostasis. In conclusion, we have shown that absence of TRPM7 channel activity strongly diminishes activation-dependent T-cell signaling, NFATc1-translocation, *IL-2* expression and secretion, as well as proliferation in both Jurkat T cells and primary human CD4 lymphocytes. Many of these effects are rescued by supplementation with MgCl_2_. Thus, TRPM7 could be a valuable pharmacological target modulating T-cell function.

## Discussion

Lymphocyte activation, specifically of T lymphocytes, is an important process with implications for the whole immune system. The ability to pharmacologically influence and reduce T-cell activation is a primary therapeutic strategy for many autoimmune defects (Walker, 2022; Sakaguchi et al., 2020; Rock et al., 2011). Therefore, further insight into the complex activation process of these cells is needed to unravel the pathogenesis and treatment options for a multitude of immunopathologies. We, here, conducted the first functional study on TRPM7 activity in primary human T lymphocytes. While TRPM7 had already been linked to numerous aspects of T-cell activation in different mouse models and cell lines (Beesetty et al., 2018; Romagnani et al., 2017; Mellott et al., 2020), we now characterize TRPM7 as an important and potentially druggable player of human lymphocyte activation. We utilized pharmacological inhibitors to study the role of TRPM7 in primary human T cells. The risk of unspecific pharmacologic effects was mitigated by validating our approach in lymphocytes in comparison to a genetic TRPM7 knockout model in Jurkat cells, and by using two different specific TRPM7 inhibitors in key experiments. Rescue experiments by supplementation with MgCl_2_ further underline the importance of TRPM7 activity for CD4 T cell function. Which proteins facilitate cellular Mg^2+^ uptake, and whether TRPM7 is one of them, has been a contentious issue in the past (Li et al., 2011; Stangherlin & O’Neill, 2018; Castiglioni et al., 2023). MagT1, long believed to be a Mg^2+^ transporter, has now been shown to be a subdomain of the N-linked glycosylation apparatus (Ravell et al., 2020). Moreover, the authors showed no alterations in total and ionized serum magnesium levels in patients diagnosed with XMEN disease, who carry a loss of function mutation in MagT1 (Ravell et al., 2020). For now the predominant interpretation seems to be TRPM7 being connected to cellular and systemic Mg^2+^ homeostasis (Zou et al., 2019; Schmitz et al., 2003; Ryazanova et al., 2004). Similar to many other cell types (Chubanov et al., 2024; Schmitz et al., 2003; Hoeger et al., 2023; Hardy et al., 2023; Mellott et al., 2020), our study further supports a role for TRPM7 as the primary Mg^2+^ uptake pathway in lymphocytes. Given that many effects of impaired TRPM7 function can be restored with Mg^2+^ supplementation, also supported by the data shown here, TRPM7-independent pathways of Mg^2+^ uptake must exist, for example through transporter proteins. Different potential Mg^2+^ transporters, such as CNNM2 and SLC41A1-3, have been proposed, but findings have so far been inconclusive (Bai et al., 2021; Mellott et al., 2020).

Recently, Mendu et al. showed mice harboring a thymus-specific deletion of TRPM7 to be resistant to Concanavalin-A-induced autoimmune hepatitis (Mendu et al., 2020). In their study, Mendu et al. reported TRPM7-deleted CD4 T cells to prefer T_reg_ lineage and non-T_reg_ CD4 cells to activate normally (Mendu et al., 2020). Partially in line with these findings, our results suggest that inhibition of TRPM7 influences iT_reg_ differentiation of human CD4 T cells, as we observed enhanced FOXP3 expression upon TRPM7 blockade. Our findings, in conjunction with the data shown by Mendu et al, highlight a possible therapeutic effect of TRPM7 inhibition in T-cell mediated autoimmune diseases. Importantly, immunological self-tolerance is mediated via naturally occurring CD4 regulatory T cells. Furthermore, these cells have been shown to play key roles in maintaining immune homeostasis, development of autoimmune diseases or graft-*versus*-host disease in patients with organ transplants (Sakaguchi et al., 2020; Haxhinasto et al., 2008; He et al., 2024). Induction of iT_regs_ is dependent on retinoic acid, short-chain fatty acids and TGF-ß. Previous findings support the notion that TRPM7 kinase moiety is influencex by TRPM7 channel conductance, while the kinase activity is not essential for channel function (Hoeger et al., 2023; Nadolni et al., 2020; Romagnani et al., 2017; Ryazanova et al., 2004). Since TRPM7 kinase has been shown to influence T-cell activation ( (Beesetty et al., 2018; Romagnani et al., 2017), this mechanism of connected channel and kinase function might very well be the case for some of the effects observed in this study and will remain subject of further investigations. However, despite several available TRPM7 channel blockers, the scientific community still lacks pharmacological tools to target TRPM7 kinase, making it especially challenging to interpret the actions of TRPM7 kinase *versus* channel function.

Activation of the AKT signaling pathway can impair T_reg_ development *in vivo*, while inhibition of this pathway, combined with TCR signaling, can induce FOXP3 expression in these cells (Sakaguchi et al., 2020; Sauer et al., 2008; Haxhinasto et al., 2008). In addition, SMAD proteins have been reported to have diverse functions in T-cell differentiation. While SMAD4 is indispensable for Th17 differentiation, deletion of SMAD2 has been suggested to promote FOXP3 transcription (Dong, 2021; Martinez et al., 2010). Of note, a direct phosphorylation of AKT SMAD2, via the TRPM7 kinase, influencing downstream signaling has recently been demonstrated for murine and human immune cells (Hoeger et al., 2023; Romagnani et al., 2017; Nadolni et al., 2020). Consequently, kinase-deficient murine naïve T cells were unable to differentiate into the pathogenic Th17 linage, while T_reg_ development was not impaired. Moreover, lack of TRPM7 kinase activity in a murine GvHD model ameliorated disease onset and severity (Romagnani et al., 2017). In line with this study, we here demonstrated for the first time that the impact of TRPM7 on pro-and anti-inflammatory T-cell homeostasis may be translated from mice to men.

Contrary to our findings showing diminished activation of human CD4 T cells after blockade of TRPM7, Mendu et al. showed that murine non-T_reg_ CD4 cells can still be activated (Mendu et al., 2020). This discrepancy could be due to functional differences in human and murine cells. Moreover, their genetic model may induce altered thymocyte development and differentiation, which is not easily comparable to physiologically differentiated cell populations. In line with our recent findings, Faouzi et al. and Beesetty et al. described TRPM7 to been linked to altered SOCE in DT40 chicken B cells and a TRPM7 kinase-deficient mouse model, respectively (Beesetty et al., 2018; Faouzi et al., 2017). However, underlying key mechanisms still remain unclear and demand further investigation.

In summary, TRPM7 is an important regulator of human T lymphocyte function regarding not only immune system homeostasis, but potentially also lymphatic malignancy. Being an important pathway for Mg^2+^ entering the cells, TRPM7 regulates T-cell signaling by influencing Mg^2+^ dependent cellular activation processes. While further research into TRPM7 and its effects on immune cell function including TRPM7 kinase related signaling is needed, this study underlines TRPM7 as a potentially druggable target in T-cell-dependent pathologies.

## Materials and Methods

### Jurkat cells and cell culture

*TRPM7*-deficient (clone E12, *KO1* and clone A03, *KO2*, both ThermoFisher Scientific) Jurkat clones were generated by CRISPR/Cas-9 genome editing at ThermoFisher Scientific (US). Primary lymphocytes and Jurkat cells (Jurkat E6.1 (WT)) were cultured in Roswell Park Memorial Institute (RPMI) medium containing 10% HI-FBS and 1% penicillin/ streptomycin in a humidified atmosphere at 37°C containing 5% CO_2_. Medium of *KO* cells was supplemented with 6 mM MgCl_2_.

### Primary human T cell isolation

Cells were isolated from peripheral blood of healthy donors according to the respective ethics approvals. PBMCs were isolated by density gradient centrifugation using Lymphoprep (Stemcell Technologies, Vancouver, BC, Canada). Isolation of respective lymphocyte subsets was achieved using magnetic cell specific separation kits. For naive CD4 T cells EasySep™ Human Naïve CD4 T Cell Isolation Kit II was used, for CD4 T cells, the EasySep™ Human CD4T Cell Isolation Kit was used. For both CD4^+^ CD25^-^ effector cells and CD4^+^ CD25^+^ T_reg_ cells, EasySep™ Human CD4^+^CD127lowCD25^+^ Regulatory T Cell Isolation Kit was used, according to the manual. A minimum of two different donors were used in primary human T cell experiments.

### TRPM7 inhibitors

Synthetic TRPM7 inhibitor NS8395 was purchased from Alomone.

Waixenicin A is a natural compound inhibitor and was isolated as following: Freeze-drive biomass of *Sarcothelia edmonsoni* Verill, 1928 was ground and extracted with hexane. After removal of solvent and elution through a C18 solid phase extraction column, the extract was subjected to reversed phase HPLC (column: SiliCycle dt C18, 30 × 100 mm, 5μm; mobile phase: acetonitrile/water gradient, 50-80% acetonitrile from 0-2 min, 80-100% acetonitrile from 2-6 min; 100% acetonitrile from 6-12 min). Waixenicin A eluted at 6,01 min and was aliquoted into 50 μg single use vials. Purity was confirmed at >95% by LC-MS with evaporative-light scattering detector.

### Electrophysiology

TRPM7 currents were acquired *via* whole-cell patch clamp. A ramp from −100 mV to + 100 mV over 50 ms acquired at 0,5 Hz and a holding potential of 0 mV was applied. Inward and outward current amplitudes were extracted at −80 and + 80 mV, respectively. Data were normalized to the cell size measured after whole-cell break-in (pA/pF). Capacitance was measured using the capacitance cancellation (EPC-10, HEKA). Mg^2+^-free extracellular solution (in mM): 140 NaCl, 3 CaCl_2_, 2.8 KCl, 10 HEPES-NaOH, 11 glucose (pH 7.2, 290-300 mOsm/l). Intracellular solution (in mM): 120 Cs-glutamate, 8 NaCl, 10 Cs-EGTA, 5 EDTA (pH 7.2, 290-300 mOsm/l).

### Proliferation and viability measurements

Jurkat cells were seeded at a density of 500,000 cells into 24-well plates and cultured in normal RPMI or RPMI with 6 mM MgCl_2_ for 5 days. Proliferation was analyzed daily using Guava ViaCount on a Guava Easycyte 12HT flow cytometer (Cytek Bioscoences, Fermont, TX, USA). Proliferation experiments on primary T cells followed a similar procedure. Alternatively, T cells were stained with CFSE dye (1 μM, Biozym), washed and cultured for 5 days, before monitoring proliferation traces (dye dilutions) on a BC Cytoflex flow cytometer.

### Inductively coupled plasma mass spectrometry

Mg^2+^ content was determined by inductive couple plasma mass spectrometry (ICP-MS) by ALS Scandinavia (Sweden). Jurkat WT and KO cells were incubated overnight in RPMI ± 6 mM MgCl_2_, washed 2x with dPBS (w/o Mg^2+^ or Ca^2+^; Sigma Aldrich). Likewise, Jurkat WT cells were cultured overnight in RPMI ± 6 mM MgCl_2_ containing 30 µM NS8593. Cells were seeded with a density of 5 million cells per condition, cell pellets were dried overnight at 70°C and stored at −80°C. Collected samples were shipped on dry ice for further analysis via ICP-MS.

### Jurkat cell Ca^2+^ imaging

Jurkat cells were loaded with 3 µM Fura-2 AM and 0.05% Pluronic^®^F-127 (Invitrogen) in imaging buffer, 15 min at 37°C. Cells were washed with imaging buffer to remove excess dye. Imaging buffer consisted of Ca^2+^ - and Mg^2+^-free HBSS supplemented with (in mM): 2 CaCl_2_, 0.4 MgCl_2_, 1 glucose. Cells were seeded into Poly-L-lysine pre-coated µ-Slide 8-well high, chambered coverslips and incubated for 10 min before start of the measurement. Time lapse images were acquired on an AnglerFish imaging system (Next Generation Fluorescence Imaging/NGFI, Graz, Austria), using 5 µM thapsigargin (Thermo Fisher) to mobilize Ca^2+^ from intracellular stores. The specific TRPM7 channel inhibitor NS8593 was used at a concentration of 30 µM. Viable cells, identified by their ionomycin response at the end of the measurement, were analyzed with Fiji.

### Ca^2+^ imaging of primary lymphocytes

Primary CD4 cells were loaded with 3 µM Fura-2 AM in RPMI supplemented with 10% FBS, 30 min at 37°C while in reaction tubes. Cells were washed twice with imaging buffer to remove excess dye. Imaging buffer contained (in mM): 140 NaCl, 2 CaCl_2_, 1 MgCl_2_, 2.8 KCl, 10 HEPES-NaOH, 11 glucose (pH 7.2, 290-300 mOsm/l). Cells were incubated for 15 min at RT and then slowly pipetted onto chambered, antibody-coated coverslips. Intracellular Ca^2+^ was monitored with Fura-2 AM (SantaCruz) using dual excitation at 340 nm and 380 nm, detection at 520 nm. Fluorescence images were acquired on a TillVisIon imaging system (TILL photonics).

### Immunofluorescence staining

Localization of NFATc1 was acquired on a Zeiss LSM 780 microscope or Zeiss LSM 900 confocal microscope, using a 63x oil objective. Jurkat cells were stimulated with 5 µM thapsigargin for 30 min or left unstimulated. Primary human T cells were stimulated with plate-bound α-CD3/α-CD28 antibodies for 45 min. TRPM7 channels were inhibited using 30 µM NS8593 and compared against cells treated with DMSO as solvent control. Cells were permeabilized with 0.1% Triton X-100 for 5 min and stained for intracellular NFAT using anti-NFATc1 antibody (1:100, Santa Cruz, #7A6) in 0.2% BSA/1% normal goat serum in PBS, and secondary anti-mouse antibody AF647 (1:1000, Cell Signaling). Cells were counterstained with DAPI (0.2 µg/mL) and mounted onto glass coverslips using Antifade ROTIMount FluorCare (Carl Roth). Zen 3.5 software was applied. Nuclear NFAT levels were analyzed, therefore regions of interest (ROI) were defined by nuclear outlines (DAPI signals). AF647 signal intensity was corrected by background signals.

### Flow cytometry of activation markers

Lymphocytes were seeded in 96-well plates at 2*10^5^ cells per condition in 100µl RPMI with 10% FBS. Cells were treated with 0.1% DMSO, NS8593 (30 µM, 20 µM or 10 µM, as indicated) or 6 mM MgCl_2_ as indicated. 15 min after treatment, cells were stimulated with antibodies against CD3/CD28 (2 µg/mL CD3 and 1 µg/mL CD28 antibodies, ImmunoCult™ Human CD3/CD28 T Cell Activator, Stemcell Technologies, or eBioscience) or PMA (20 ng/mL and ionomycin (1 µg/mL) (both from SigmaAldrich). After 24 or 48 h, respectively, cells were stained according to the manufacturer’s instructions. Cells were washed twice after staining. Isotype controls or FMO controls were performed. Cells were analyzed using a Guava Easycyte 6-2L flow cytometer (Luminex Corporation, Austin, TX, USA), or a Beckman Coulter CytoFLEX. The following antibodies were used: anti-human CD4-VioBlue (Miltenyi REA623), anti-human CD45RA-APC-Vio770 (Miltenyi, REA562), anti-human CD69-APC (Miltenyi, REA824), anti-human CD25-VioBright515 (Miltenyi, REA570).

### IL-2 quantification

Lymphocytes were seeded in 96-well plates at 2*10^5^ cells per conditions in 100 µl RPMI with 10% FBS. Cells were treated with 0.1% DMSO, 30 µM NS8593, or 6 mM MgCl_2_ as indicated. 15 min after treatment, cells were stimulated with antibodies against CD3/CD28 (ImmunoCult™ Human CD3/CD28 T Cell Activator, Stemcell Technologies, as before). Cell supernatants were collected 48 h after cell stimulation and stored at −80°C. IL-2 concentrations were analyzed using a Biogems Precoated Human IL-2 ELISA kit (Biogems International, Inc., USA) according to manufacturer’s instructions by measuring absorbance at 405 nm on a BMG Labtech Clariostar Plus plate reader.

### mRNA isolation

Jurkat TRPM7 KO cells were cultured overnight in normal RPMI without additional MgCl_2_ supplementation, KO cells and WT cells were seeded at a density of 4*10^6^ cells per conditions and stimulated for 3 h with 10 ng/µL PHA. mRNA was isolated from cell pellets using RNeasy Mini Kit (Qiagen) following manufacturer’s instructions. mRNA concentrations were determined via OD measurement.

### cDNA synthesis and quantitative real-time PCR (qRT-PCR)

For cDNA synthesis, 0.5 µg mRNA was diluted in H_2_O, mixed with 1 mM dNTPs (Promega) and 0.5 µg Oligo(dT)_12-18_ (Promega) and incubated for 5 min at 70°C. On ice, 5x First-Stand Buffer, SuperScriptTM II Reverse Transcriptase (Promega) and DEPC-treated H_2_O was added and incubated for 60 min at 42°C. The resulting cDNA was diluted 1:4. Transcripts were analyzed by specific primer pairs. Master mixes additionally contained cDNA and SYBR-Green^TM^ (Sigma-Aldrich). Transcripts were measured in technical triplicates on a CFX-96 cycler (BioRad): 50°C 2’, 95°C 10’ (preincubation), 95°C 15’’, 62°C 30’’, 72°C 30’’, 40 cycles (amplification), 95°C 10’’, 60°C 1’ (melting), 40°C 10’ (cooling). Primer pairs (all human 5′-3′): hIL2 (fw) TTTACATGCCCAAGAAGGCC and (rev) GTTGTTTCAGATCCCTTTAGTTCCA and hHPRT1 (fw) CCCTGGCGTCGTGATTAGTG and (rev) TCGAGCAAGACGTTCAGTCC. A minimum of three independent experiments were performed. CT values of housekeeping transcripts were subtracted from measured CT values, to calculate 2^(−ΔCT) values.

### iTreg differentiation and flow cytometry staining

Naïve CD4 T cells were seeded at a density of 1* 10^5^ cells per condition into a 96-well plate, and treated with 30 µM NS8593 or equivalent volume of DMSO. Induction medium contained a-CD3/a-CD28 dynabeads (ThermoFisher), 10 ng/µL rh IL-2 (Immunotools), 5 ng/µL TGF-ß (Immunotools) and 100 nM ATRA (Sigma Aldrich). Cells were cultured for 6 days in a humidified atmosphere at 37°C containing 5% CO_2_, with intermediary medium exchange on day 4. Cells were analyzed using a Guava Easycyte 6-2L flow cytometer (Luminex Corporation, Austin, TX, USA). The following antibodies were used: anti-human CD4-VioBlue (Miltenyi REA623), anti-human CD25-PE (BioLegend, BC96), anti-human CD45RA-APC-Vio770 (Miltenyi, REA562), anti-human CTLA4-BV605 (BioLegend, BNI3), anti-human FoxP3-APC (Miltenyi, REA1253). Naïve CD4 T cells were used as gating control.

### Ethics

Peripheral blood of healthy volunteers was obtained by venipuncture. The study was conducted according to the guidelines of the Declaration of Helsinki and, approved by the local ethics boards of the Johannes Kepler University Linz (EK 1064/2022) as well as the Ludwig-Maximilians-Universität München (Az.21-1288).

### Statistics

Data were plotted using Graphpad Prism 8 (Graphpad Software, Boston, MA, USA) or higher. Statistical analysis of the difference of two data sets was performed using Student’s T-test or Mann Whitney U test. Comparison of three or more data sets was performed using one- or two-way-ANOVA, Kruskal-Wallis test or Friedmann test, depending on the respective experimental design.

## Supplementary Materials

Supplementary information is available in the online version of the manuscript.

Supplementary Figure 1: Validation of Jurkat TRPM7 KO clone 2 shows reduced proliferation and activation

Supplementary Figure 2: Apamin as control substance for potential off target effects on NS8593

Supplementary Figure 3: T cell isolation controls and additional FACS data

Supplementary Figure 4: Dose-response curve of TRPM7 inhibitor NS8593 on CD4 T cells

## Acknowledgments

We thank Viktoria Sperrer for her excellent technical assistance. Authors thank the following funding agencies: KH was supported by the FoeFoLe program (LMU Munich); SZ was supported by the Deutsche Forschungsgemeinschaft (DFG, German Research Foundation) TRR 152 Project 14 (SZ), 15 (TG) and 16 (AD). FDH received support from NIH NIGMS P20GM103466.

## Author contribution

KH, AM, BH and SZ wrote the manuscript. KH, AM and BH performed main experiments. DL, MW, TH and MD performed additional experiments. FDH isolated and purified a natural product inhibitor. MS, AD and TG provided valuable expertise and feedback. SZ conceived and supervised the study. All authors revised the manuscript and agreed on publishing.

## Data and material availability

Materials may be requested from the corresponding author.

## Abbreviations

AUC: area under the curve
ICP-MS: inductively coupled plasma mass spectrometry
IL-2: interleukin 2
KO: knock out
NFAT: nuclear factor of activated T cells
SOCE: store-operated Ca2+ entry
TCR: T cell receptor
Treg: regulatory T cells
TRPM7: Melastatin-like Transient Receptor Potential, member 7
WT: wild type

---

**Supplementary Figure 1:**
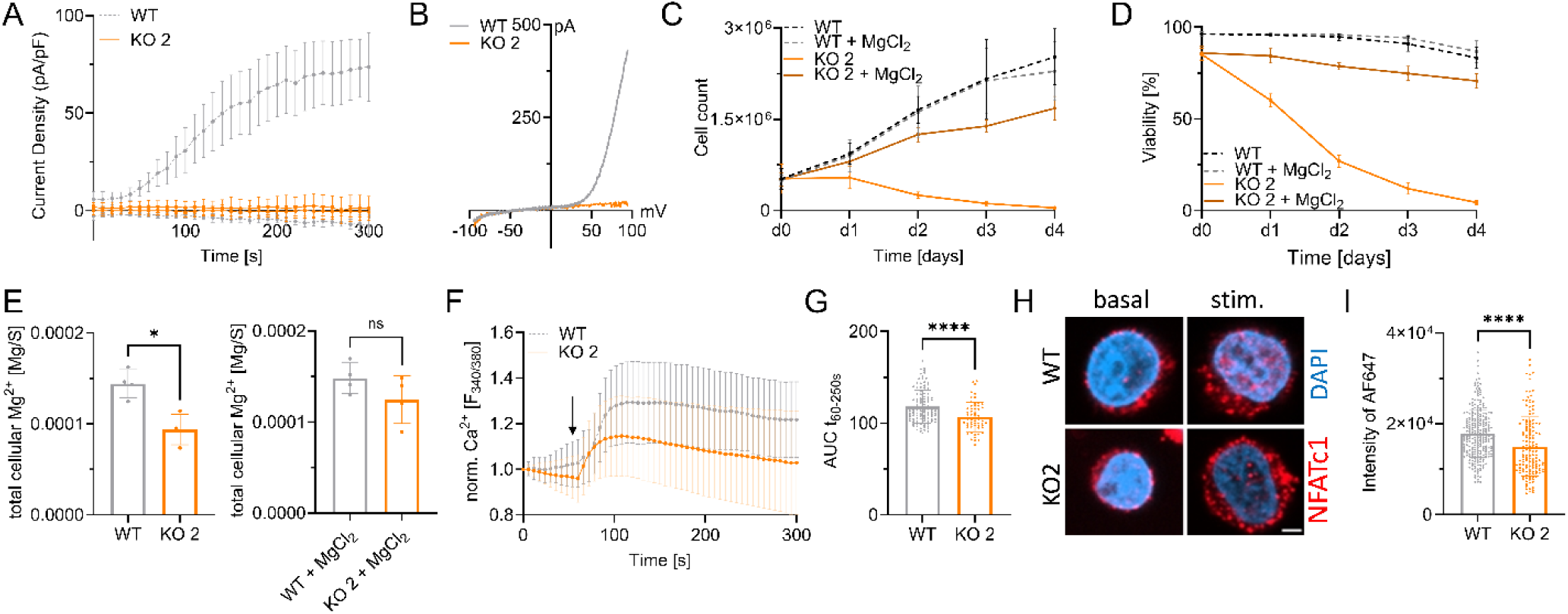
Validation of Jurkat TRPM7 KO clone 2 shows reduced proliferation and activation. A) TRPM7 current densities and B) TRPM7 I/V relationship of Jurkat cells during whole-cell patch clamp experiment with Mg^2+^-free intracellular solution. WT (WT, grey) and TRPM7 KO2 Jurkat clone (KO2, orange). n(WT)=9; n(KO2)=10. C) Cell counts and D) viability of natively proliferating TRPM7 WT and KO2 Jurkat clone in RPMI medium with 10% FBS, with and without supplementation with 6 mM MgCl_2_. n=3, measured in duplicates. E) Cellular Mg contents quantified by ICP-MS. WT and TRPM7 KO2 Jurkat clone, cultured in regular (WT-)media or in medium supplemented with 6 mM MgCl_2_ for 18 h ahead of sampling, n=4. F) Fura-2 based imaging of cytosolic Ca^2+^ concentration of Jurkat cells. Stimulation with 5 µM thapsigargin at indicated time point (arrow). WT (WT, grey) and TRPM7 KO2 (KO2, orange) Jurkat clone, n (WT) =111; n (KO2) = 59; G) Quantification of the area under the curve (AUC) of respective curves shown in F. H) Representative immuno-fluorescent images of NFATc1 localization in WT and KO2 clone before (basal) and after 30 min stimulation (stim.) with 5 µM thapsigargin, scale bar = 2 μm. NFATc1 in red, DAPI in blue. I) Quantification of nuclear NFATc1 levels upon stimulation of TRPM7 WT (WT, grey) and KO (KO2, orange) clone. n (WT) = 261; n (KO2) = 149. Statistics: Two-way ANOVA (C, D), one-way ANOVA (E) or Student’s t test (G, I). * P<0.05; and **** P<0.0001. Data are mean ± SD.

**Supplementary Figure 2:**
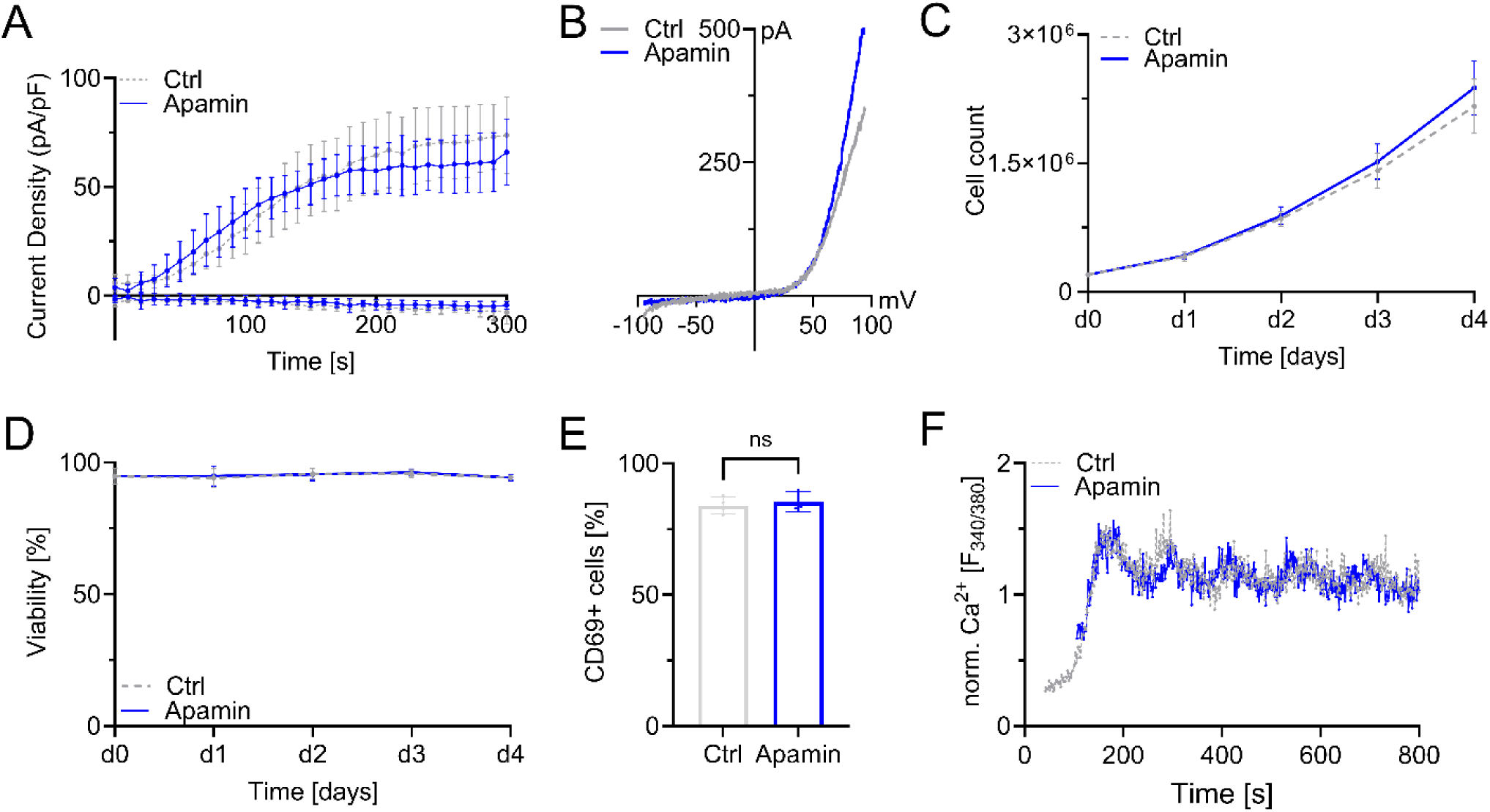
Apamin as control substance for potential off target effects of NS8593. A) TRPM7 current densities and B) TRPM7 I/V relationship of Jurkat T cells during whole-cell patch clamp experiment with Mg^2+^-free intracellular solution. Controls (Ctrl, grey) and cells treated with 1 µM apamin (Apamin, blue), n (Ctrl)=9, n (Apamin)=6. C) Cell counts and D) viability of natively proliferating Jurkat cells in RPMI medium with 10% FBS, with and without 1 µM apamin (Apamin, blue), n=4. E) Flow cytometry of upregulation of activation markers CD69 in primary CD4 T-lymphocytes 48 h after anti-CD3/CD28 stimulation. Cells treated either as control (Ctrl, grey) or with 1 µM apamin (Apamin, blue), n=4. F) Representative trace of CD4 T cells Fura-2 based imaging of cytosolic Ca^2+^ concentrations following anti-CD3/CD28 stimulation. Antibodies bound to microscopy chamber bottom with cells sinking down in saline containing 2 mM Ca^2+^ during running measurement, coming to rest in focus plane with contact to stimulation antibodies. Cells measured as control (Ctrl, grey) or in presence of 1 µM apamin (Apamin, blue). Statistics: Student’s t test (D). n.s.—not significant. Data are mean ± SD.

**Supplementary Figure 3:**
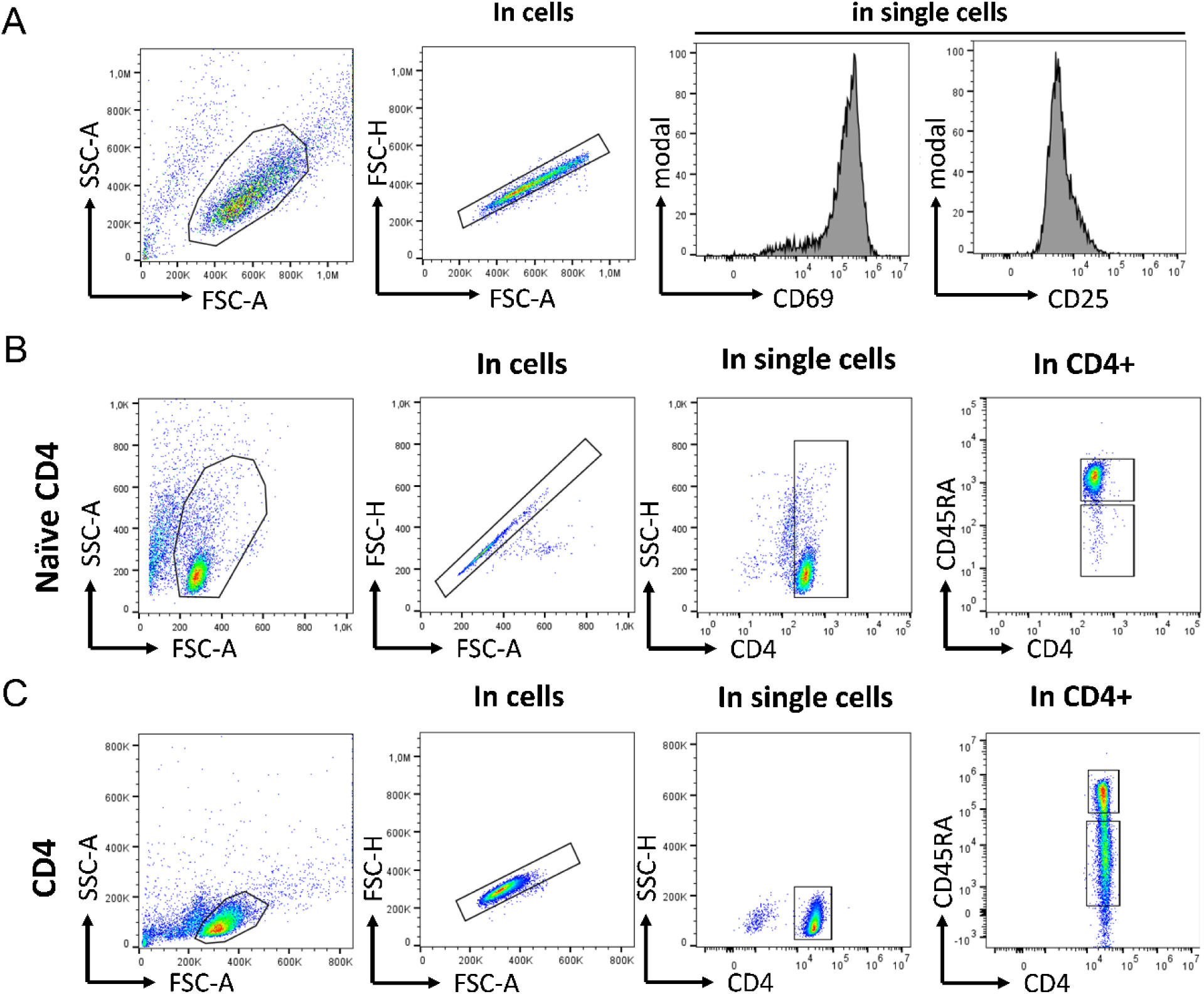
T cell isolation controls and additional FACS data. A) Representative FACS plots and gating strategy for CD69 and CD25 visualization, shown for Jurkat WT cells. B) Representative FACS plots and gating strategy to confirm identify of isolated naïve CD4 T cells and C) conventional CD4 T cells.

**Supplementary Figure 4:**
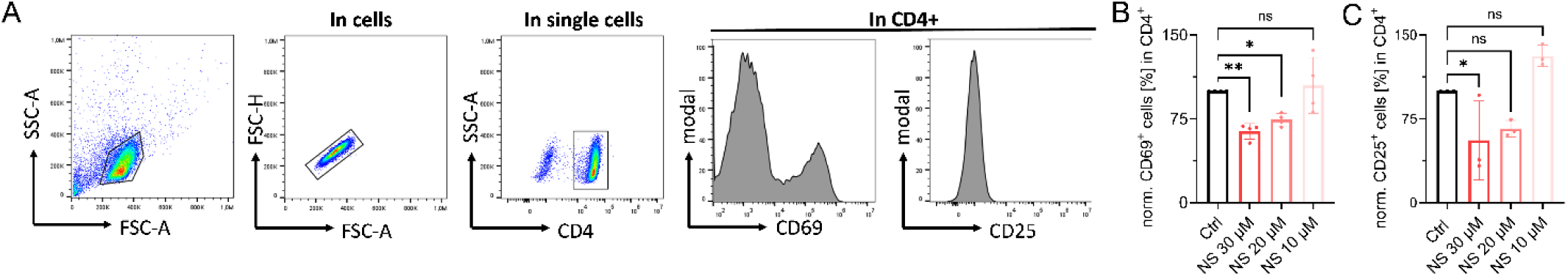
Dose response curve of TRPM7 inhibitor NS8593 on CD4 T-cell activation. A) Representative FACS plots and gating strategy for CD69 and CD25 shown for conventional CD4 T cells. B+C) Quantification of flow cytometry data of NS8593 dose-dependent upregulation of CD69 (B) and CD25 (C) expression on conventional CD4 T cells, 48 h after anti-CD3/CD28 stimulation or PMA/ionomycin stimulation, respectively, n=3-4. Statistics: One-way ANOVA (B, C). * P<0.05; ** P<0.005 and n.s.—not significant. Data are mean ± SD.

